# Synapses from the high-order thalamic nucleus and motor cortex are co-clustered in the distal tuft dendrites of the mouse somatosensory cortex

**DOI:** 10.1101/2020.11.08.363200

**Authors:** Nari Kim, Sangkyu Bahn, Joon Ho Choi, Jinseop S. Kim, Jong-Cheol Rah

## Abstract

The posterior medial nucleus of the thalamus (POm) and vibrissal primary motor cortex (vM1) convey essential information to the barrel cortex (S1BF) regarding whisker position and movement. Therefore, understanding the relative spatial relationships of these two inputs is critical prerequisites to acquire insight into how S1 synthesizes information to interpret the location of an object. Using array tomography, we identified the locations of synapses from vM1 and POm on distal tuft dendrites of L5 pyramidal neurons. We found that synapses from vM1 and POm are spatially clustered on the same set of dendrites with unusually high density. Furthermore, the clusters of vM1 and POm synapses colocalize each other. These findings suggest that synaptic clusters, but not dendritic branches, act as functional units and cooperatively contribute to nonlinear dendritic responses.

## INTRODUCTION

The primary somatosensory cortex (S1) is an essential area to integrate sensory and modulatory inputs upon sensing objects. The apical dendrites of thick-tufted layer 5 pyramidal (TTL5) neurons of S1 are branched into dense oblique dendrites in L1 and receive various synaptic inputs. These include the feedback information associated with high-level cognitive functions such as attention and modulation of plasticity (Audette et al., 2019; Gambino et al., 2014; Gilbert and Sigman, 2007; Sjöström and Häusser, 2006). The primary sources of such information include a high-order thalamic nucleus, posterior medial nucleus (POm), and vibrissal primary motor cortex (vM1) (Deschênes et al., 1998; El-Boustani et al., 2020; Kinnischtzke et al., 2014; Petreanu et al., 2009; Veinante and Deschênes, 2003).

POm neurons integrate sensory signals from the brain stem (Alloway et al., 2003) and feedback inputs from various cortical areas (Groh et al., 2014; Veinante et al., 2000) to encode the information about whisker position and movement (Diamond et al., 2008; Yu et al., 2006). On the other hand, vM1 appears to deliver signals related to muscle output change, such as initiation and delicate control of whisking movement (Carvell et al., 1996; Diamond et al., 2008; Ferezou et al., 2007; Sreenivasan et al., 2016), to the barrel field of S1 (S1BF). Therefore, it is plausible to hypothesize that the integration of the inputs from POm and vM1 in S1 is critical for decoding an object touch in the context of whisker location (O’Connor et al., 2010).Then, in which structural level would the two synaptic inputs be integrated?

The axons from POm and vM1 are highly ramified and spatially intermingled in L1 of the S1BF (Deschênes et al., 1998; Koralek et al., 1988; Lee et al., 2013; Veinante and Deschênes, 2003; Wimmer et al., 2010). These inputs densely innervate distal tuft dendrites of L5 pyramidal neurons (Petreanu et al., 2009; Zhang and Bruno, 2019). However, for synaptic inputs received on distal dendrites to propagate down to soma, strong voltage attenuation by passive dendritic filtering has to be overcome (Magee and Cook, 2000; Rail, 2018; Williams and Stuart, 2002). To this end, regenerative dendritic activity is essential. To support this idea, Larkum and colleagues discovered N-methyl-D-Aspartate (NMDA) receptor channel-dependent spikes in distal tuft dendrites that are critical to evoke calcium spikes and thereby action potentials, as well as response selectivity (Larkum et al., 2009; Smith et al., 2013). However, the NMDA spikes are usually confined within the dendritic branch as they become greatly attenuated at the bifurcation point (Larkum et al., 2009; Palmer et al., 2014). Therefore, it is suggested that each synaptic input is clustered together on the same set of branches to overcome this limitation and that dendritic branches serve as a functional unit of neural networks (Branco and Häusser, 2010). Regarding L5 pyramidal neurons, the two synaptic inputs (POm and vM1) could either be spatially clustered in the same branch or clustered into different branches (“dendritic tropism”)(Polsky et al., 2004).

Another level of synaptic integration can be suggested by recent findings showing that POm inputs can enhance the efficiency of sensory signaling in the S1BF. Activation of POm inputs evokes reliable NMDA receptor-mediated dendritic spikes and mediates rhythmic whisking-derived synaptic potentiation in L2/3 neurons (Gambino et al., 2014). Paired activation of POm and feedforward inputs enhance the feedforward transmission by disinhibiting vasoactive intestinal peptide (VIP) – expressing neurons (Williams and Holtmaat, 2019). Enhanced and prolonged whisking-evoked cortical activity by POm activation was reported in L5 neurons as well (Audette et al., 2019; Mease et al., 2016). Therefore, essential synaptic inputs might be localized to the proximity of POm synapses for efficient integration.

To test these hypotheses, (1) spatial clustering of synaptic inputs, either on the same branch or different branches and (2) co-clustering of vM1 near POm synapses, we analyzed the relative distribution of the vM1 and POm synapses using array tomography (AT). AT is a high-resolution, wide-field fluorescence microscopy technique that achieves isotropic resolution by repeated imaging of ultrathin serial sections followed by computational volumetric reconstruction (Micheva and Smith, 2007). Because AT provides ~200 nm isotropic resolution, it allows synapse detection with an accuracy as high as ~80% (Micheva and Smith, 2007; Mishchenko, 2010; Rah et al., 2013). Fluorescent labeling of multiple molecules or input sources is easily achievable (Mishchenko, 2010; Rah et al., 2013), allowing a distinction between inputs from vM1 and POm as well as dendrites from TTL5 neurons. Furthermore, the imaging time to reconstruct a significant volume of neurons is remarkably shorter than electron microscopy (Micheva and Smith, 2007; Mishchenko, 2010; Rah et al., 2013).

Using AT, we imaged the synaptic inputs from vM1 and POm on distal dendrites of TTL5. We observed extremely dense innervation of the two synaptic inputs on an overlapping set of dendritic branches. Synapses showed a robust input-wise synaptic clustering, suggesting that the synaptic clusters, not dendritic branches nor individual synapses, may function as a fundamental unit of integration. Also, we observed the colocalization of vM1 clusters to the proximity of POm clusters, implying a cooperative interaction of the two types of inputs.

## RESULTS

To examine the relative distribution of synapses from vM1 and POm on the distal dendrites of L5 pyramidal neurons, we reconstructed 18 dendritic branches and measured the locations of the two types of synapses with array tomography (AT) (Figure 1, Materials and Methods). As previously described (Deschênes et al., 1998; Veinante and Deschênes, 2003), both axons from POm and vM1 densely innervated L1 of the S1BF (Figure 1A). Synapses from vM1 and POm were unambiguously resolved in AT (Figure 2B and C) (Collman et al., 2015; Oberti et al., 2011; Rah et al., 2013). The annotated synapses from vM1 and POm on distal tuft dendrites were quantified (Figure 1D and 2, Materials and Methods). Approximately twice as many vM1 synapses were found as POm synapses (1,566 vM1 synapses and 783 POm synapses in a total of 1626.2 μm of the reconstructed dendrites).

**Figure 1.**
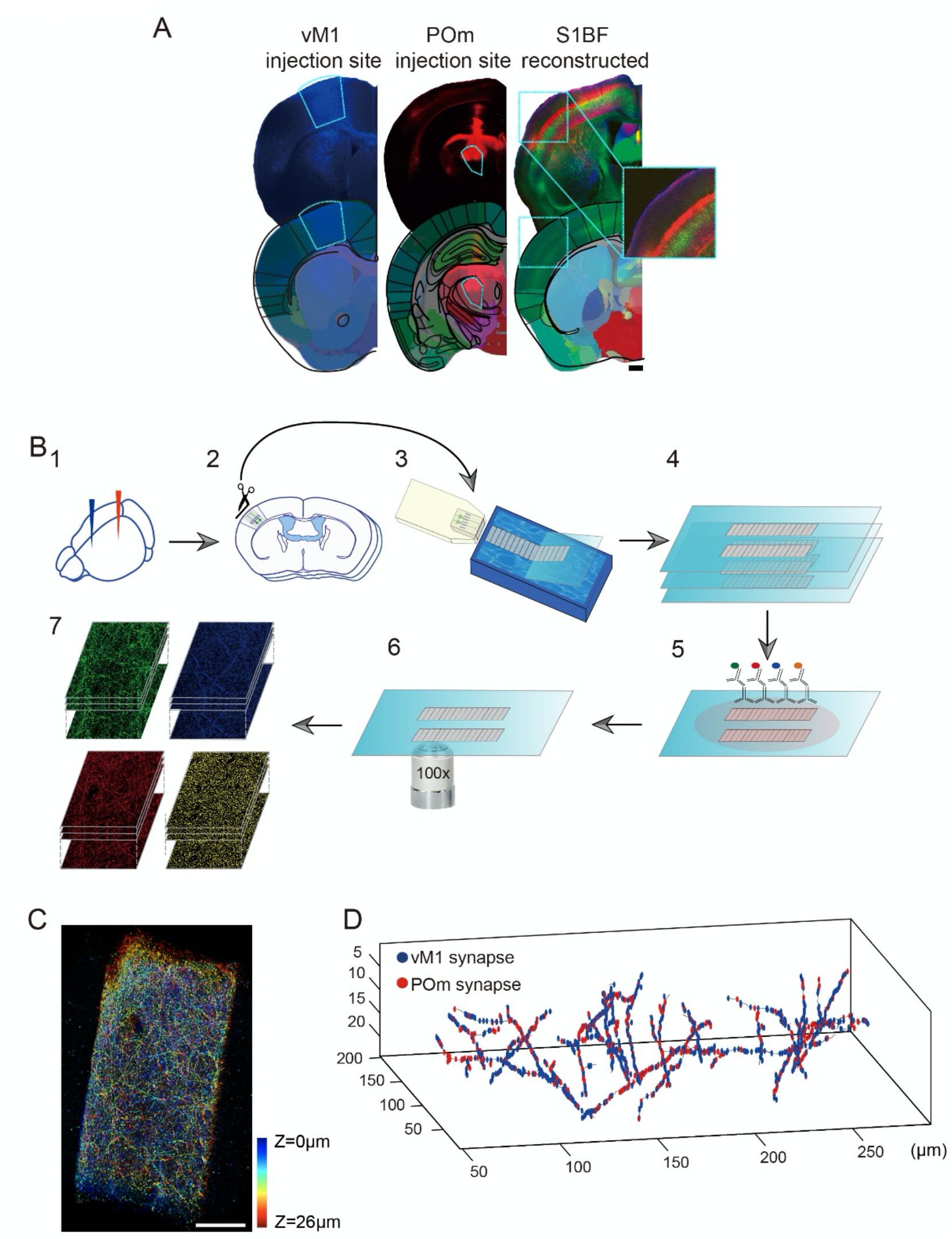
Schematic diagram of the study. (A) vM1 neurons and POm neurons were labeled with myc (left, blue) and tdTomato (middle, red) for the identification of axons from the origins. Thick tufted L5 pyramidal neurons were identified in Thy1-YFP (type H). Injection sites were validated by the pattern of innervated axons (upper) and by aligning the acquired images with the Allen Brain Reference Atlas with QuickNII (lower). The target injection sites (vM1 and POm) and reconstructed S1BF were marked with dotted white boxes. Scale bar= 500 μm. (B) Diagrammatic representation of array tomography. S1BF with vM1 and POm axons labeled were dissected for tissue embedding (B_1_-B_2_). Serial sections were acquired from embedded tissue (B_3_-B_4_). Images on the serial sections were acquired with an automated image focusing system (B_6_) after immunostaining of synaptic markers (B_5_). Four-colored, 2×3 image tiles were acquired for subsequent reconstruction (B_7_). (C) Reconstructed distal dendrites of TTL5. Scale bar = 50 μm. (D) vM1 and POm synapses were manually annotated in the reconstructed distal tuft dendrites of L5 pyramidal neurons. Blue and red circles denote synapses from vM1 and POm, respectively.

**Figure 2.**
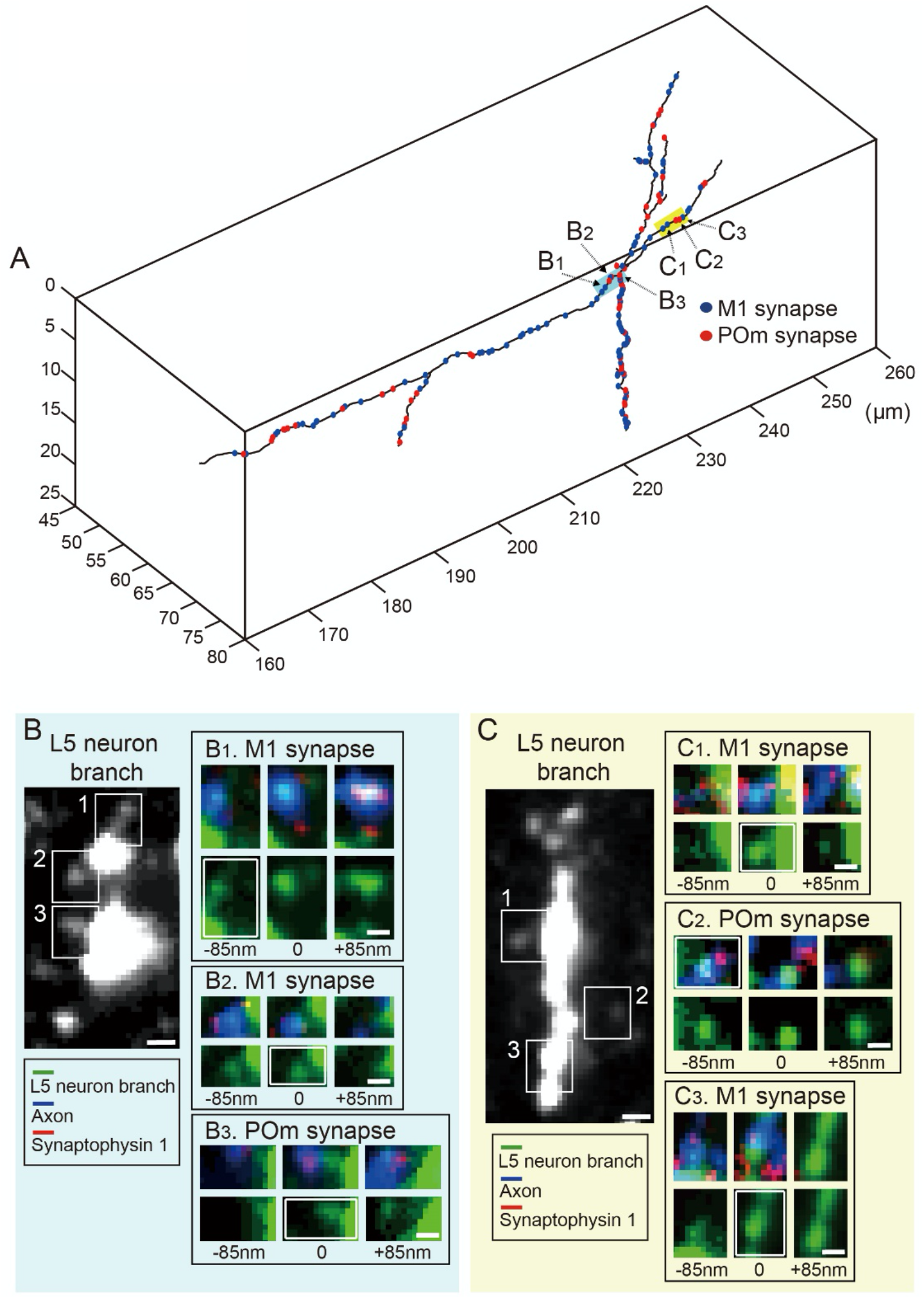
Synapses from vM1 and POm were unambiguously identified with AT. (A) A fraction of the reconstructed distal tuft dendrite with annotated vM1 (blue) and POm (red) synapses. (B) Representative image of YFP-expressing dendrite (left panel) with three examples of synapses from vM1 and POm identified by AT (B_1_-B_3_). (C) Another representative image of YFP-expressing dendrite with representative synapses (C_1_-C_3_). Green: distal tuft dendrites of L5 pyramidal neurons. Blue: axons from vM1 or POm, pseudocolored for demonstration. Red: Synaptophysin-1. Scale bar = 500 nm (left panels of B and C), 300 nm (insets).

### Synapses from vM1 and POm impinge on an overlapping set of distal dendrites of pyramidal neurons

We first examined if the integration of POm and vM1 occurs at the level of individual branches. We examined the relationship between the number of synapses and the length of dendritic branchlets, where a branchlet is the segment of dendritic branch separated by branching points (Figure 3). If all the dendritic branchlets have an equal chance to form synapses with vM1 and POm axons, the longer dendritic branchlets will have more synapses from vM1 and POm. On the contrary, if dendritic branchlets are preferred by one input and unfavored by the other, the relationship will deviate significantly from linearity. Therefore, we counted the respective number of synapses from the two input sources on every branchlet of the 18 dendritic branches. Also, to exclude the possibility that observed results are attributed to chance, we compared the results with those from the sets of the dendritic branches with randomly shuffled synapses. These synapse-shuffled dendritic branches are repeatedly used below in other analyses as well. For brevity, we call the genuine synapse distribution “observed” and the shuffled one “randomized” throughout the manuscript. The results showed that synapses from both areas are randomly distributed along the dendritic branches, as measured by the level of correlation between the number of synapses and the length of branchlets (Figure 3A and B, Pearson’s correlation coefficient, *r* = 0.768 and 0.766 for vM1 and POm synapses, respectively). Thus, this result rejects the hypothesis that synapses from vM1 and POm are organized in a branchlet-specific manner. To further confirm, we examined the relative preference of each branchlet for the two input sources by devising a metric called the preference index (PI) (see Materials and Methods). A branchlet with more definite synaptic input preference will be assigned a value closer to either +1 or −1 (+1 for vM1 inputs and – 1 for POm inputs), whereas branchlets with no preference will have PI values close to zero. The majority of PI values of analyzed branchlets were close to zero (Figure 3C) and fell within the expected range of randomized synapses (Figure 3C shaded). The only exceptions were those branchlets with a very short length, <10 μm, with only a few synapses, where accurate preference cannot be measured.

**Figure 3.**
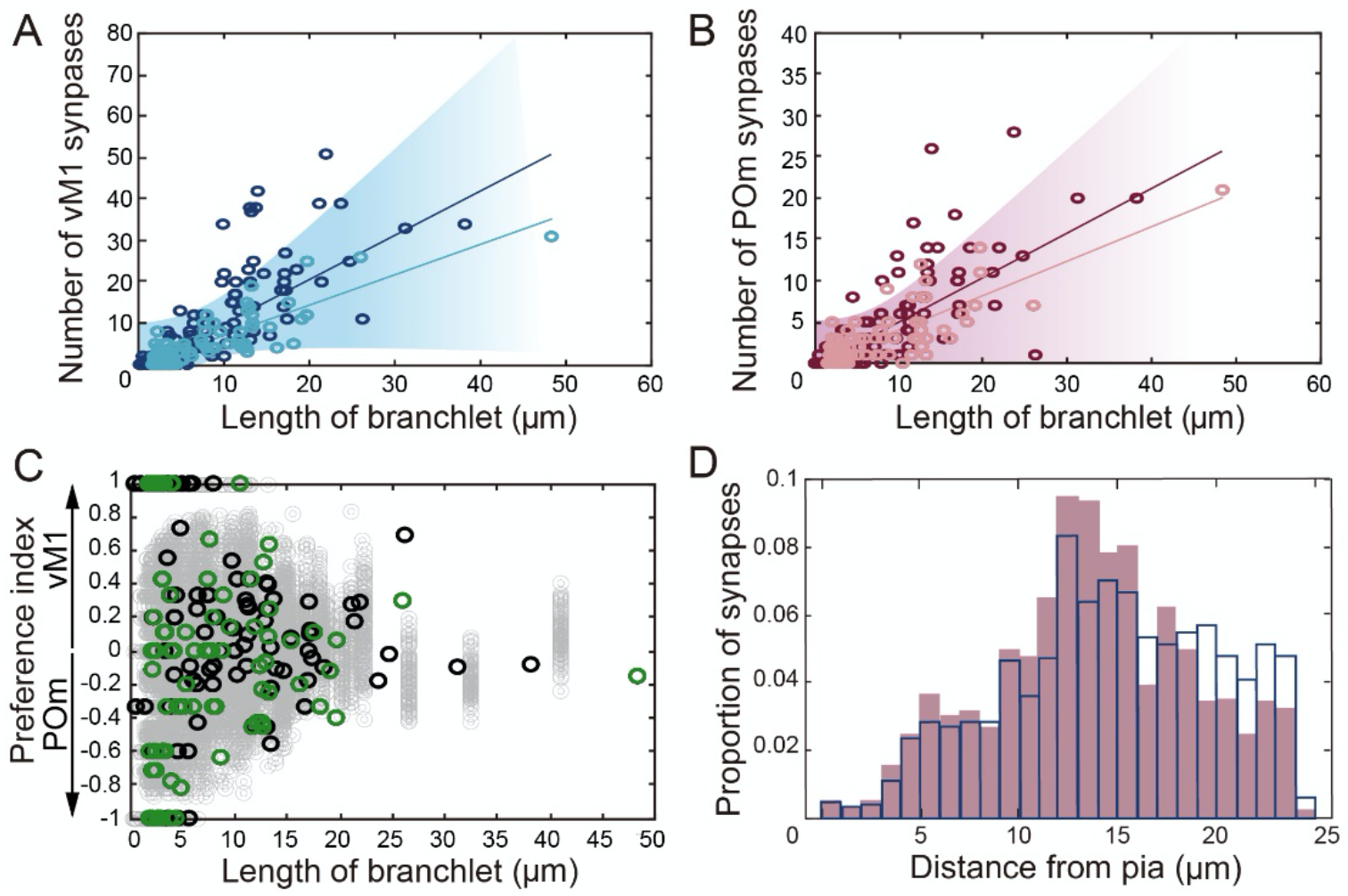
Branch tropism of vM1 and POm synapses. The number of synapses from vM1 (A) and POm (B) as a function of the length of the branchlet. The values of the end branchlets were highlighted with a lighter color. Shaded areas indicate the 95% confidence interval. (C) Preference indices (PI) of measured dendritic branchlets (black open circles). The preference indices of the end branchlets were separately denoted in green circles. The preference indices calculated from random synaptic distribution are displayed in light gray circles for comparison. (D) The distribution of synapses from vM1 (blue open bar) and POm (red filled bar) along with the distance from the pial surface.

This result was somewhat unexpected, considering the whisker-object touch specific Ca^2+^ transient occurred in a subset of distal tuft dendrites that were dependent upon vM1 activity (Xu et al., 2012). Branchlet-wise input organization could be specific for the end branchlet because segregated inputs will be integrated into the mother branchlets, which would obviate the need to have segregated inputs on the low-order branchlets. Consistent with this idea, a functional study has demonstrated dramatic functional segregation selectively at the terminal branches (Cichon and Gan, 2015). Therefore, we re-examined whether the terminal branchlets may have stronger input tropism. However, synapses on end branchlets showed approximately the same degree of linearity (Figure 3A and B, light-colored circles, Pearson’s correlation coefficient, *r* = 0.8493, 0.8140 for vM1 and POm, respectively) and PI values as the other branchlets (Figure 3C, green circles) and random distribution (Figure 3C shaded). Also, we examined if the two inputs differentially innervate based on the laminar location of dendrites (Figure 3D). We failed to detect a significant difference in density or the distance from pia between the input sources. Together with the axon distribution illustrated in Figure 1A, we concluded that dendritic branchlets are not the functional units that receive the uniformed synaptic inputs, but that the two inputs impinge on the same set of dendritic branchlets.

### Synapses from vM1 and POm are spatially clustered on distal dendrites

We then examined how the two types of synaptic inputs are organized within a dendritic branch. Functional studies have demonstrated high functional correlations between closely located synapses, suggesting the possibility that the synapses sharing the inputs can be clustered at small patches within a dendritic branch (Kleindienst et al., 2011; Makino and Malinow, 2011). Indeed, spatial clustering of synaptic inputs with the same sources has been reported in other dendritic compartments and brain areas (Druckmann et al., 2014; Gökçe et al., 2016; Rah et al., 2013). To examine whether the spatial synaptic clustering is a general feature of synaptic wiring, including in distal dendrites, we examined the pattern of distribution using three independent methods. First, we compared the distribution of intersynapse distances of the observed synapses. The distances between two successive, mutually independent events that occur randomly and constantly follow the exponential distribution. Therefore, if the inter-synapse distance fits an exponential distribution, it can be inferred that the synapses are located randomly. The inter-synapse intervals of both vM1 and POm deviated substantially from the exponential curve. In particular, the greatest deviation from the exponential relationship was observed at very short inter-synapse intervals (Figure 4A and B), suggesting a spatial clustering of synapses for both inputs. We then compared the distribution of the nearest neighbor distance for observed synapses with random distribution. The cumulative probability histogram of the nearest-neighbor distance was significantly shifted toward short distances in observed synapses (Figure 4C and D, Kolmogorov-Smirnov test, *p* = 3.31 x 10^-24^ for vM1 synapses, 1.32 x 10^-15^ for POm synapses), suggesting that the synapses with the same input sources tend to localize close together compared to the case of random distribution of synapses.

**Figure 4.**
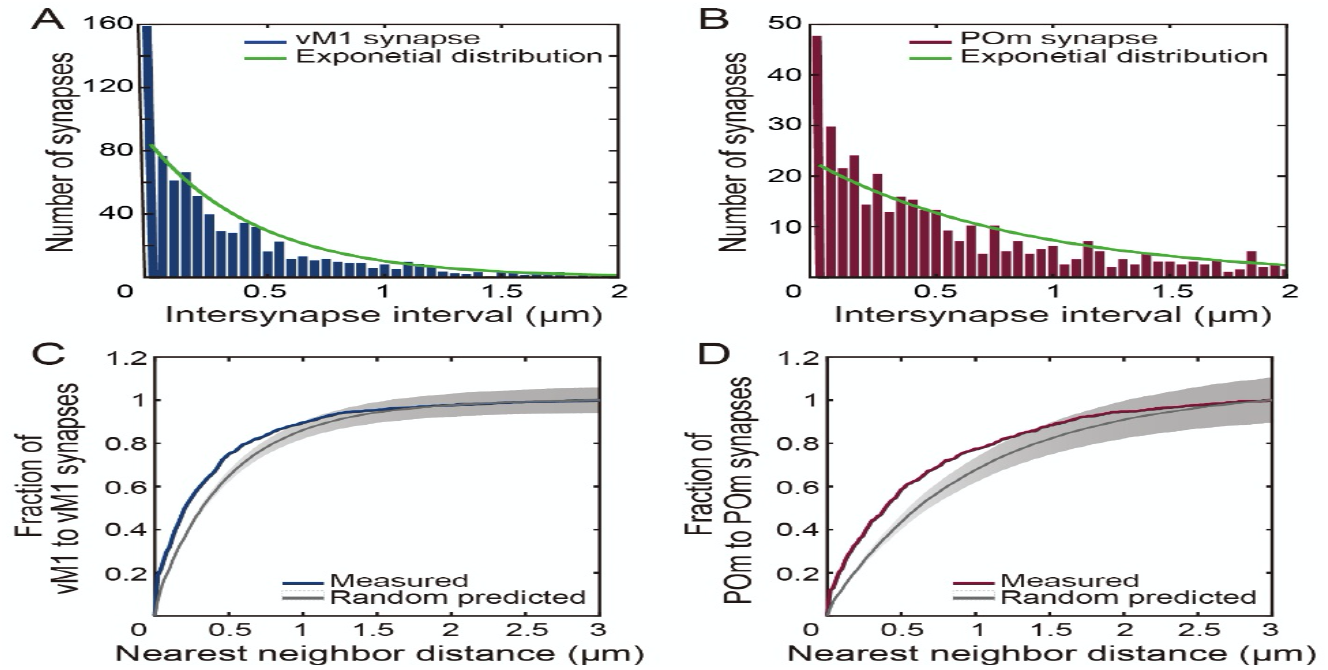
Inter-synapse intervals and nearest neighbor distance. (A, B) Histogram of inter-synapse distance among synapses from vM1 (A) and POm (B). The exponential distribution expected by random distribution is illustrated in green lines. (C, D) The nearest neighbor distance of the same types of synapses. Cumulative distribution of nearest neighbor distances between the synaptic inputs from vM1 (C) or POm (D) in comparison to the random distribution (gray, mean with 95% confidence interval).

### Synapses from vM1 and POm form spatial clustering more often and with greater magnitude than a random distribution

Consequently, we examined whether synaptic clusters are found more frequently than is expected by chance. We followed the method suggested by Yadav and colleagues with slight modifications (Yadav et al., 2012). Briefly, to determine a synaptic cluster we grouped synapses using a single-linkage hierarchical clustering algorithm with the mean distance of nearest neighbor as a dissimilarity criterion (Figure 5A and B, see Materials and Methods). As expected, the probability histogram of the number of clustered synapses from the 10,000 random shufflings exhibited roughly a Gaussian distribution (Figure 5C). To quantify the magnitude of synaptic cluster formation of a given branch, we introduced a measure dubbed the “clustering score” or C-score. The C-score of a branch was defined as the proportion of dendritic branches with fewer synaptic clusters than the genuine branch itself amongst the shuffled dendritic branches (Yadav et al., 2012). In other words, a C-score of 0.99 would mean that 99% of the randomized branches have fewer synaptic clusters than the branch. Therefore, the more likely the observed synaptic clustering can occur by chance, the closer the C-score will be to 0. With a separate set of random synaptic distribution (30 random distributions of 18 branches), we compared the distribution of the C-scores with the measured synaptic distribution. Both branches with vM1 and POm synapses showed significantly higher C-scores than is expected by random synaptic distributions (Figure 5D, median C-score of the measured and simulated vM1 synapses were 0.98 and 0.47; median C-score of the measured and simulated POm synapses were 0.90 and 0.42, respectively; One way ANOVA, **** *p* < 0.0001). We then compared the number of dendritic branches with highly clustered synapses by using the C-score threshold of 0.9, 0.95, and 0.99 (the vertical lines in Figure 5C). The dendritic branches with highly clustered synapses were observed significantly more frequently in the measured vM1 and POm synapse distribution than those in random distribution in all thresholds tested (Figure 5E and F; binomial test, * *p* < 0.5, **** *p* < 0.0001). Next, we characterized the observed synaptic clusters (Figure 6). A significantly greater proportion of synapses participated in forming synaptic clusters than simulated branches (Figure 6B-E; Wilcoxon matched-pairs signed-rank test, one-tailed, **** *p* < 0.0001 and ** *p* = 0.0069). Individual synaptic clusters consist of the higher number of synapses (Figure 6F and G; Mann Whitney test, one-tailed, **** *p* < 0.0001 and * p = 0.042) in a slightly longer (vM1) or similar (POm) length of synaptic clusters in measured branches, compared with their simulated branches (Figure 6H and I; Mann Whitney test, one-tailed, * *p* = 0.0108). We conclude that both synapses from vM1 and POm form robust spatial clustering of their own on the overlapping set of dendritic branches. Based on all the analyses above (Figures 4–6), we concluded that the synaptic inputs from vM1 and POm on distal dendrites of L5 pyramidal neurons are spatially clustered. Our result is in line with previous studies demonstrating spatially clustered distribution of thalamocortical inputs on dendrites of L5 neurons in the mouse barrel cortex (Rah et al., 2013), CA3 inputs on CA1 neurons in the hippocampus (Druckmann et al., 2014), and intralaminar inputs on the basal dendrites of L5 pyramidal neurons (Gökçe et al., 2016). Therefore, it is tempting to conclude that the spatial clustering of synapses from an identical origin can be a general wiring principle regardless of the subcellular location or brain regions.

**Figure 5.**
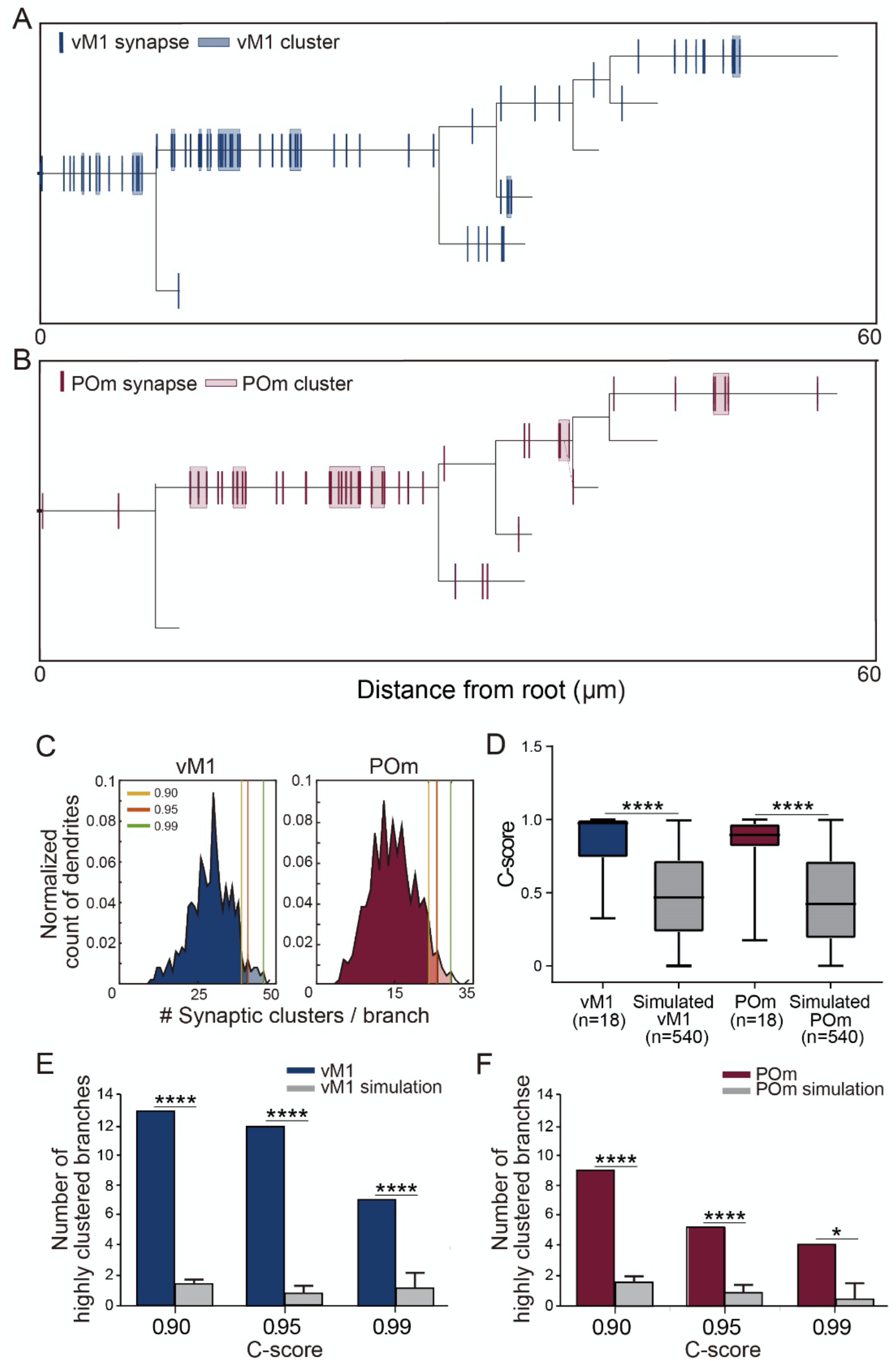
Clustering analysis of synapses with different inputs on dendritic branches. (A, B) Example dendrograms with the locations of synaptic inputs from vM1 (A) and POm (B). Groups of synapses determined as synaptic clusters were denoted with colored boxes. (C) Normalized histogram of the number of synaptic clusters generated by 1,000 random synaptic distributions on 18 reconstructed dendrites. Three thresholds to determine “highly clustered dendrites” (C-score = 0.90, 0.95, and 0.99) are denoted with the vertical colored lines. (D) C-score distribution from branches with measured synapses and separately simulated 30 random synaptic locations on 18 reconstructed dendrites (gray). (E, F) The number of branches with highly clustered synapses from vM1 (E) and POm (F) is determined with various C-scores compared with the same set of branches with 30 randomized synaptic locations (gray).

### Synapses from vM1 and POm are clustered together

Next, we examined the relative location of the two types of synaptic clusters. If the two types of clusters are on the overlapping part of the dendrites, cooperative dendritic spikes can be expected. In particular, if POm synapses function as a gain enhancer of sensory signals as proposed, close positioning of vM1 synapses would enhance the chance of successful integration. On the other hand, synaptic clusters with spatial exclusion will competitively interact with each other due to the refractory period of voltage-gated ion channels necessary for the regenerative dendritic activity (Remy et al., 2009). To understand the spatial relationship between the two types of synapses, we calculated the distance of the closest POm synapse from each vM1 synapse and *vice versa* (Figure 7A). We then compared the distribution of the distances with that of randomly allocated synapses (Figure 7B and C). To our surprise, the distance of vM1-to-POm synapses, as well as that of POm-to-vM1 synapses, was significantly outranged toward shorter distance than expected by the random allocation (Kolmogorov-Smirnov test; *p* = 3.31 x 10^-9^ for vM1-to-POm synapses, 1.49 x 10^-13^ for POm-to-vM1 synapses), suggesting that the two different inputs tend to localize close to each other. As the two types of synapses are spatially clustered, we examined the relative locations of the two types of synaptic clusters. Consistent with the nearest neighbor distance between different types of synapses (Figure 7B and C), we found that vM1 clusters are located closer to POm clusters than randomly distributed clusters (Figure 7D; Kolmogorov-Smirnov test, *p* = 0.0137).

To quantify the proximity of the two types of synapses, we further devised a new concept, called the “synaptic cloud model”. The term, synaptic cloud model, intends to make an analogy to the electron cloud model. As the location of electrons in an atom is described by a probability distribution, we regard each synapse to exist probabilistically following the Gaussian distribution centered around its actual location with 500 nm of variance, as exemplified in Figure 8A. The summation of such a distribution of a type of synapse at any location of the branch is the expected number of synapses along the branches. The higher the value for a location is, the more synapses of one type densely exist around the location. For brevity, we call the expected number of synapses of one synapse type as the “synaptic cloud density” (SCD). We then asked whether SCD for vM1 synapses (SCD_vM1_) varies along with the SCD for POm synapses (SCD_POm_). Corroborating the notion that the two types of synapses localize near each other, we found that the distribution of SCD_vM1_ was shifted upward where SCD_POm_ is high (SCD_POm_>0.35) compared to the rest of the dendrites (Figure 8A inset, and 8B; median SCD_vM1_ was 0.07 and 0.45 for low SCD_POm_ and high SCD_Pom_, respectively; Wilcoxon rank sum test, *p*=7.93 x 10^-12^). As the SCD is the sum of the Gaussian distribution centered around actual synapses, the value has to be smaller at the end of each branchlet. To control the inherent covariation of the measure, we repeated the comparison in the random distribution. Indeed, we found a slightly upper-shifted SCD_vM1_ distribution on high SCD_POm_, but the magnitude of the difference was significantly smaller (Figure 8B; median SCD_vM1_ was 0.080 for low SCD_POm_ vs. 0.10 for high SCD_POm_; Wilcoxon rank sum test, p=7.54 x 10^-6^). Finally, we directly compared the mean SCD_vM1_ values on the portion of dendrites with varying SCD_POm_ range. The mean SCD_vM1_ stayed relatively constant regardless of the SCD_POm_ up to the maximum SCD_POm_ in the random distribution (Figure 8C, gray). On the contrary, the mean SCD_vM1_ increased as the SCD_POm_ increased (Figure 8C, blue). In other words, the chance to find vM1 synapses is significantly higher in a particular location where the number of neighboring POm synapses are high (Figure 8B and C).

## DISCUSSION

Synaptic voltage deflection is attenuated and temporally smeared in a distance-dependent manner (Mainen et al., 1996; Rall, 1967; Rall et al., 1967; Stuart and Spruston, 1998; Williams and Stuart, 2002). Despite the electrotonic disadvantages, substantial excitatory synaptic inputs impinge on distal tuft dendrites where there is nearly a mm-long path distance before reaching an action potential initiation zone (Csabai et al., 2018; Santuy et al., 2018). To overcome the electrotonic disadvantage by the location and to preserve the information against passive dendritic filtering, nature designed various ways of compensation. Relatively high impedance and low capacitance because of the morphology of distal dendrites, as well as the contribution of differential ion channel distribution, counterbalance the filtering to some degree (Bernander et al., 1994; Jaffe and Carnevale, 1999; Magee, 1999; Rall, 1967). An increase in the amplitude of excitatory postsynaptic potentials (EPSPs) with distance from the soma has been reported to compensate for the disadvantage in hippocampal pyramidal neurons (Magee and Cook, 2000; Vilas Menon et al., 2013). However, the effect of the distance-dependent synaptic scaling is somewhat limited, especially in the neocortex (London M, 2001; Williams and Stuart, 2002).

We suggest that a large quantity of intense excitatory synaptic inputs on distal dendrites may ensure successful information transfer to mitigate the electrotonic disadvantages of the distal synaptic inputs. Synapses from vM1 and POm are as densely packed as ~1 and 0.5 synapses per μm of dendrites, respectively. The densities of the synapses are exceedingly high compared to the mean spine density of the TTL5 neurons that is estimated between 1 and 2 spines per μm (Larkman, 1991). The excitatory synaptic density on distal tuft dendrites has never been studied thoroughly, but reportedly, ~10% of entire dendritic spines are located on distal tuft dendrites of thick-tufted L5 pyramidal neurons (Larkman, 1991; Rubio-Garrido et al., 2009). The reported density could have been underestimated, considering the estimation was made using optical imaging with an anisotropic resolution where spines perpendicular to the focal plane are obscured by the dendritic shaft (Feldman and Peters, 1979; Larkman, 1991).

Array tomography (AT) used in the current study is an appropriate technique to answer this question because the resolution along the Z-axis is not determined by Abbe’s rule but by the thickness of the serial sections, ~85 nm. With the improved Z-axis resolution to 200 nm, previous studies have revealed that the synapse detection accuracy is as high as 80% compared to that by transmission electron microscopy (Mishchenko, 2010; Rah et al., 2013). Although the current study lacks the reexamination of the accuracy of synapse detection in distal dendrites, we believe that the current study provides at least a comparable accuracy with the previous estimation. The size of excitatory synapses on distal tuft dendrites are comparable with the synapses used to estimate the synapse detection accuracy in previous studies (Santuy et al., 2018). Furthermore, to secure better Z-axis resolution, we used even thinner serial sections (85 nm) than the previous study. Therefore, it is unlikely that the detection accuracy is lower than the previous estimations. This argument is supported by the fact that the number of excitatory synapses on distal dendrites is in a similar range of previous studies using electron microscopy (Csabai et al., 2018).

Dendritic spikes boost the contribution of synaptic inputs to the somatic membrane potential and, thereby, AP generation (Larkum et al., 2009; Major et al., 2013). The generation of dendritic spikes critically relies on the spatiotemporal cooperativity among synaptic inputs to spatially defined dendritic compartments (Branco and Häusser, 2010; Gasparini et al., 2004; Gasparini and Magee, 2006; Losonczy and Magee, 2006; Schiller et al., 2000; Wei et al., 2001). In particular, the generation of dendritic spikes at distal dendrites is further compartmentalized due to multiple factors. The primary source of the dendritic spike in distal tuft dendrites is NMDA receptors, which require depolarization and glutamate to be activated (Larkum et al., 2009; Palmer et al., 2014). Distal dendrites have a high density of the HCN channels and A-type K^+^ channels, which will reduce the time window for dendritic integration and decrease the chance to generate dendritic spikes (Atkinson and Williams, 2009; Berger et al., 2001; Harnett et al., 2013; Hoffman et al., 1997; Kole et al., 2006; Larkum et al., 2009). Back-propagating AP, which causes Ca^2+^ influx and reduces the threshold for a dendritic spike, is restricted in distal dendrites due to the cable properties (Brunner and Szabadics, 2016; Larkum et al., 1999).

We, therefore, examined if distal dendrites contain a substantial level of spatial clustering and an exceedingly high density of synapses, presumably to ensure a successful generation of dendritic spikes. We utilized a series of independent methods to quantify the synaptic clustering. Consistently across these methods, we observed that substantial spatial clustering of the synapses with the same origin tends to form clusters. Namely, the distance between the pairs of all synapses was closer (Figure 4A and B), and the nearest neighbor distance was significantly shorter (Figure 4C and D). Furthermore, we determined synaptic clusters with a single-linkage algorithm with the average nearest neighbor distance as the dissimilarity criterion. Near 50% of synapses appeared to be populated in less than 10% of the dendritic portion, and the length of each synaptic cluster was in the range of a micrometer, where ~4 synapses were populated (Figure 6). However, we found no dendrite-wise preference of the synapses (Figure 3).

**Figure 6.**
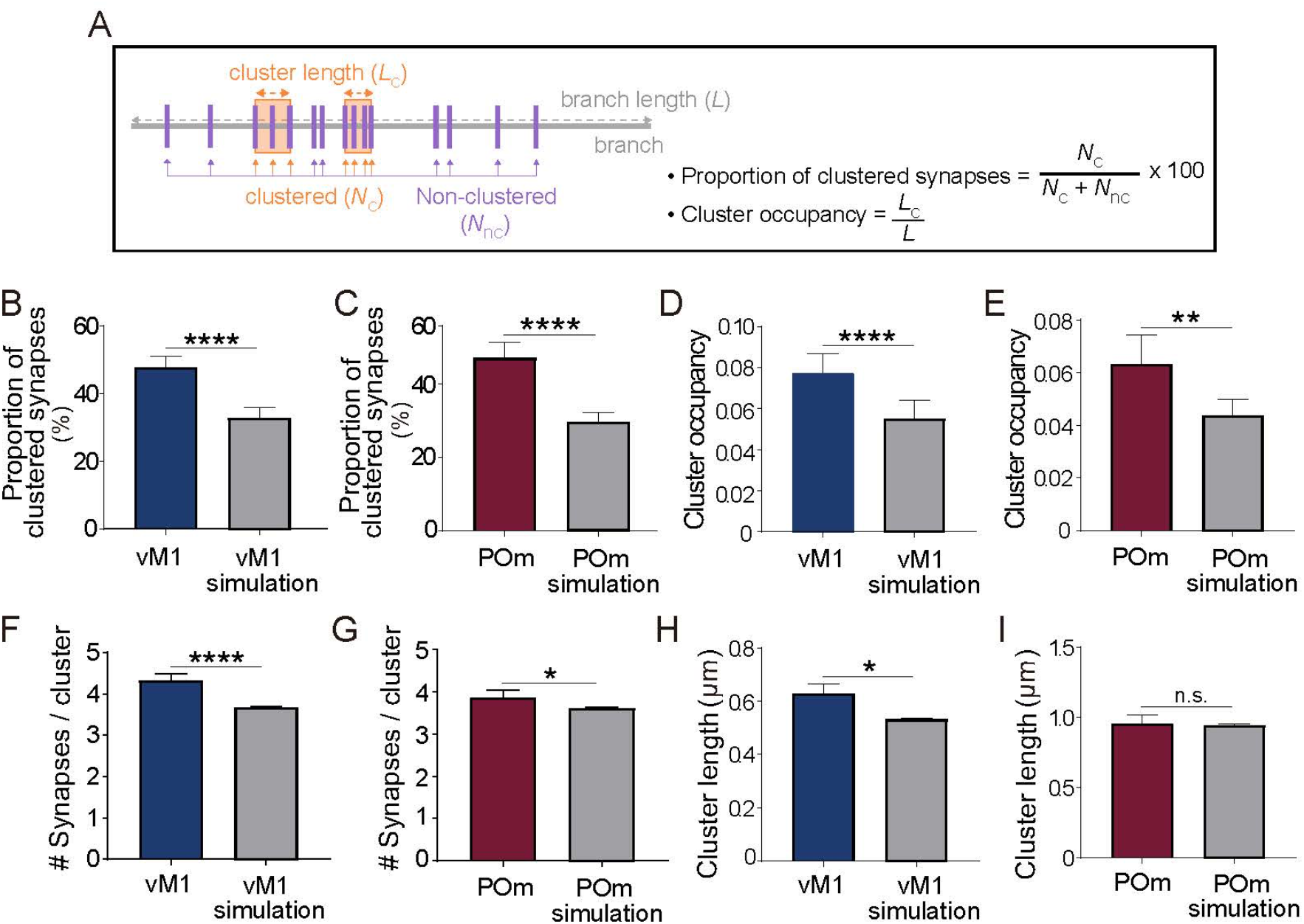
Characterization of synaptic clusters. (A) Schematic illustration to describe the terms used for the analysis. (B, C) The fraction of synapses participating in synaptic clusters of vM1 (B) and POm (C). (D, E) The cluster occupancy, the integrated cluster length divided by the length of the entire dendritic branches of vM1 (D) and POm (E). (F, G) The mean number of synapses in a vM1 (F) and a POm cluster (G). (H, I) The integrated cluster length for vM1 (H) and POm (I) synapses. Randomized simulation results are shown in gray (B to I).

Our result is reminiscent of a previous functional study that demonstrated local Ca^2+^ transients observed in 1-5 μm active dendritic subregions in response to sensory stimuli (Jia et al., 2010). We believe that the observed Ca^2+^ transients on discrete dendritic segments could be the result of spatially clustered synaptic transmission. Furthermore, the previous study found that the Ca^2+^ responses have an orientation selectivity and that the receptive fields of the responses were heterogeneous even within a dendritic branch. Our observation of no branch-wise organization of synapses may correspond with this finding (Figure 3). Together, our results provide the anatomical groundwork to suggest that neither individual synapse nor the dendritic branch is the fundamental functional compartment, but clusters of synapses function as the electrical processing units in distal dendrites of pyramidal neurons in the neocortex.

We found that the clusters of vM1 synapses and POm synapses are closely located to each other (Figures 7 and 8). We believe that synaptic clusters of the two inputs can cooperatively contribute to the nonlinear dendritic responses.

**Figure 7.**
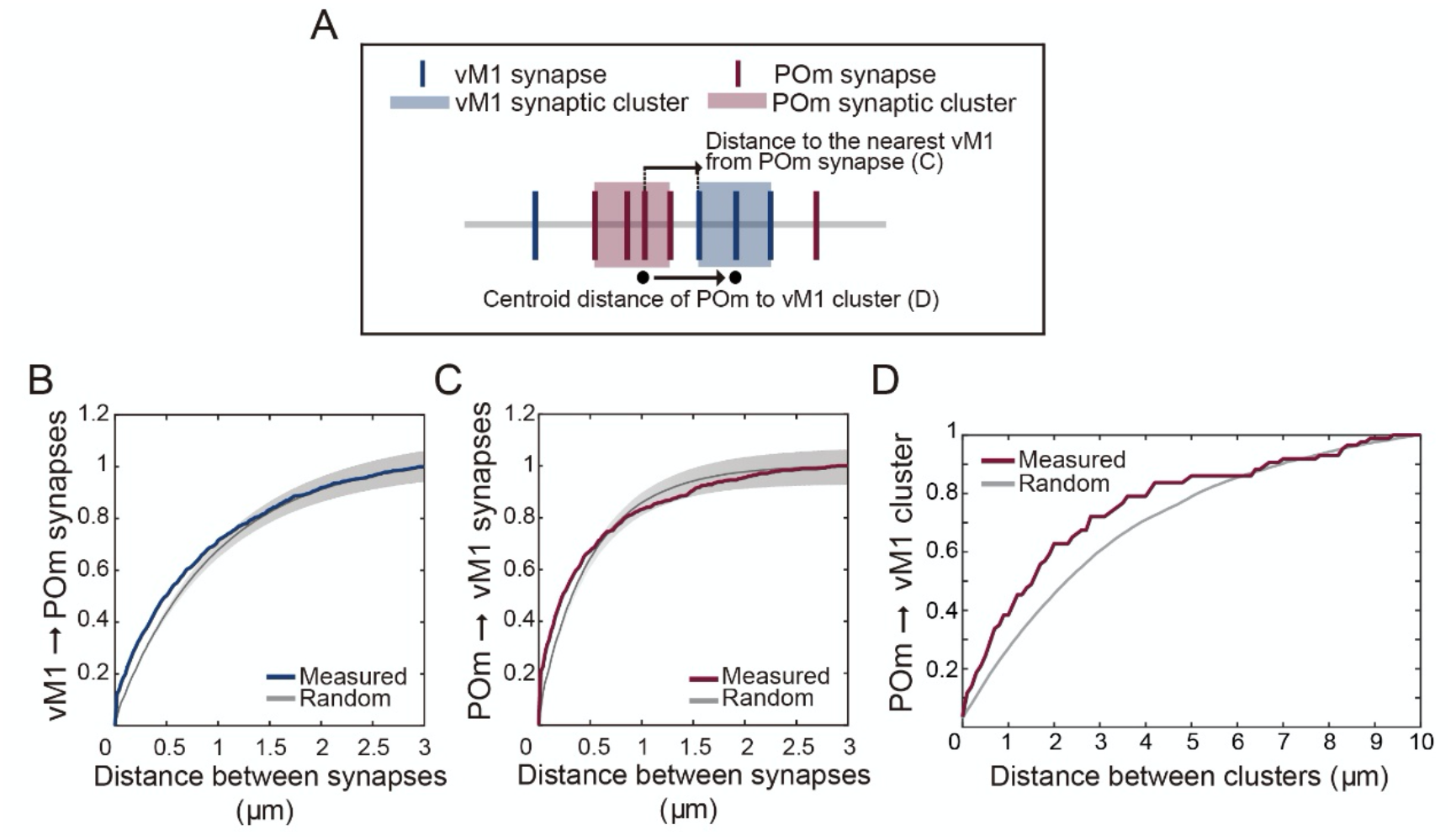
The spatial distribution between synapses with different origins. (A) Schematic illustration to describe the terms used for the analysis. (B, C) Cumulative histogram of distances to the nearest POm synapse from individual vM1 synapses (B) and distances to the nearest vM1 synapses from POm (C). Gray lines with shaded areas indicate the 95% confidence interval from simulated branches. (D) Cumulative histogram of distances to centroids of the nearest vM1 synaptic cluster from the centroid of the POm synaptic cluster (red) compared with the distances between randomized locations of synaptic clusters (gray).

**Figure 8.**
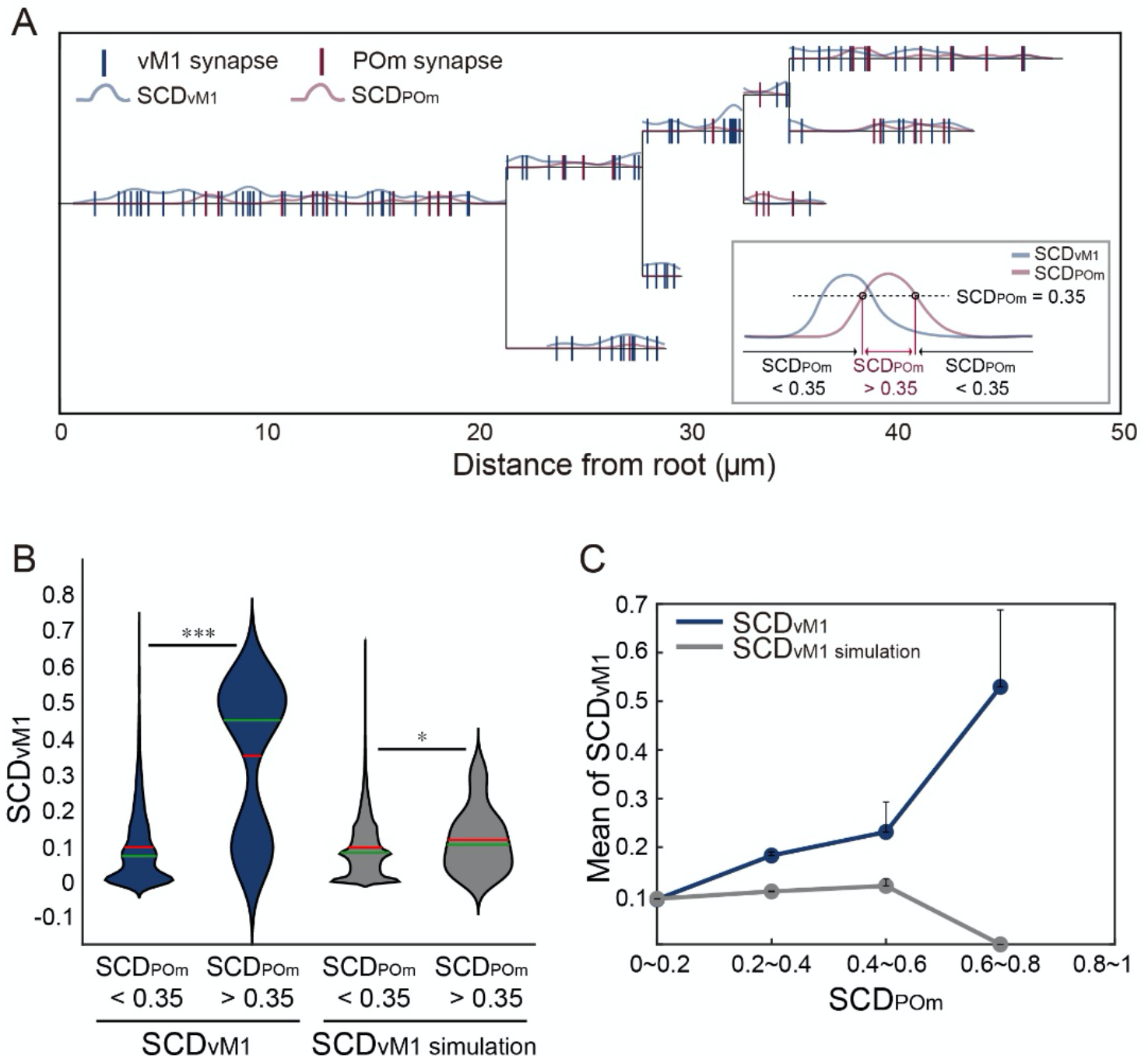
Covariation of the synaptic cloud density for vM1 (SCD_vM1_) with the synaptic cloud density for POm (SCD_POm_) (A) An example dendrogram with an SCD_vM1_ and SCD_POm_ and locations of synapses. Blue and red vertical lines denote the locations of synapses from vM1 and POm, respectively. The SCDs for both types of synapses, calculated as the sum of Gaussian distribution centered around each synapse, are illustrated along the dendrite (see METHODS). (A-inset) Diagrammatic illustration of distinction of the dendritic portion with high and low SCD_POm_. SCD_POm_ 0.35 was used for distinction criteria. (B) Distribution of SCD_vM1_ on the dendritic fraction with high and low SCD_POm_ compared with simulated random distribution, with 0.35 as a discrimination point. Horizontal red and green lines are the mean and median of each distribution, respectively. (C) The mean SCD_vM1_ at a fraction of dendrites with varying SCD_POm_ from measured (blue) and simulated random (gray) synaptic distribution.

Synaptic inputs from POm have been shown to evoke strong NMDA receptor-dependent subthreshold depolarization on dendrites in L1 of the barrel cortex in response to repetitive whisker deflection at the frequency range of natural whisking (Gambino et al., 2014). Furthermore, optogenetic activation of POm caused delayed and persistent depolarization of cortical neurons (Zhang and Bruno, 2019). Therefore, from the perspective of vM1 synapses, locations near POm synapses are advantageous to evoke reliable dendritic spikes.

We propose that POm inputs and vM1 inputs may function as an “AND” logic gate to evoke dendritic spikes upon an active object touch. A previous study by Xu et al. described vM1 activity and active touch-dependent dendritic Ca^2+^ responses on distal tuft dendrites (Xu et al., 2012). The dendritic response required active whisker movement and object touch. Neither passive whisker touch nor free whisking evoked as strong Ca^2+^ responses (Helmchen et al., 1999; Varga et al., 2011; Xu et al., 2012). Considering vM1 delivers signals related to muscle output change such as initiation of whisking rather than whisking itself (Carvell et al., 1996; Sreenivasan et al., 2016), the active object touch-specific dendritic responses could be the result of vM1-mediated responses induced by touch-evoked sudden initiation of whisking on top of the whisking-evoked prolonged depolarization by POm synapses (Ahlssar et al., 2000; Diamond et al., 1992; Ferezou et al., 2007; Jouhanneau et al., 2014; Masri et al., 2008; Matyas et al., 2010; Xu et al., 2012; Yu et al., 2006; Zhang and Bruno, 2019).

The current study presents a model in which clusters of vM1 and POm synapses colocalize and cooperatively synthesize the active touch-dependent Ca^2+^ responses in the distal dendrites. Functional studies examining how the two inputs interact in the generation of dendritic spikes or dendritic spike modulation in the context of animal behavior will be a future study of interest.

## MATERIALS AND METHODS

### Animals

All the animal experiments followed procedures approved by the Korea Brain Research Institute (KBRI). A 16-week-old Thy1-YFP (type H) male mouse (Jackson Laboratory, 003782) was used for the current study (Feng et al., 2000).

### Stereotaxic delivery of adeno-associated virus

Inputs from vM1 and POm were labeled by stereotaxic delivery of adeno-associated virus (AAV) transducing tdTomato and GFPsm-Myc (Viswanathan et al., 2015) in a Thy1-YFP (type H) mouse, respectively (Figure 1A and B_1_). TTL5 neurons were sparsely labeled in the mouse line (Feng et al., 2000). 100 nl of AAV1-CAG-tdTomato-WPRE-SV40 (Addgene, 105554) was injected in POm (AP −2.06, ML 1.25, DV 2.97 mm) and the same volume of AAV1-CAG-GFPsm-Myc-WPRE-SV40 (Addgene, 98926) was delivered to the vM1 region (AP 1.34, ML 1.50, DV 0.35 and 0.75 mm). After four weeks, the mice were sacrificed for AT. The correct injection was examined in two ways. We examined the injection sites with reference to the Allen Brain Reference Atlas using QuickNII (Figure 1A lower) (Puchades et al., 2019). Also, the pattern of axonal distribution in cortical layers was compared with previous studies (Figure 1A upper) (Deschênes et al., 1998; Petreanu et al., 2009; Veinante and Deschênes, 2003).

### Brain sampling, resin embedding, and sectioning

Four weeks after stereotaxic virus injection, the mice were deeply anesthetized with isoflurane and transcardially perfused with 4% paraformaldehyde (Electron Microscopy Sciences, 15710) in Trisbuffered saline (TBS) (VWR Life Science, K859). After further incubation in fixatives overnight, the extracted brains were rinsed and sectioned coronally at 300 μm with a vibratome (Leica, VT100S) (Figure 1B_2_). S1BF with proper labeling was dissected to embed in HM20 (Electron Microscopy Sciences, 14340) in a low-temperature embedding system (Leica, EM AFS2). The embedded tissue with a block face size of 300 μm × 400 μm was serially sectioned at 85 nm with an ultramicrotome (RMC products, PowerTome PC) (Figure 1B_3_). 297 consecutive sections were collected on coverslips (Electron Microscopy Sciences, 3423) for subsequent imaging (Figure 1B_4_).

### Immunostaining

Sections were incubated with 50 mM glycine (Sigma, G7403) at room temperature for 5 minutes to remove free aldehyde groups. The sections were permeabilized with 0.05% Triton X-100 (Fisher scientific, BP151-500) and blocked with 0.1% bovine serum albumin (BSA) (Sigma, A7030) and 0.05% Tween-20 (Sigma, P9416) for 10 minutes at room temperature. The sections were then incubated with the following primary antibodies (Figure 1B_5_): anti-GFP antibody 1:1000 (Abcam, ab13970), anti-synaptophysin-1 1:200 (Synaptic system, 101-004), anti-RFP 1:100 (MBL International, PM005), and anti-Myc 1:100 (Millipore, 05-419), and subsequently with the fluorophore-conjugated secondary antibodies (Figure 1B_5_): Alexa Fluor® 488 AffiniPure Donkey Anti-Chicken 1:000 (Jackson ImmunoResearch, 703-545-155), DyLight™ 405 AffiniPure Donkey Anti-Guinea pig IgG (H+L) 1:500 (Jackson ImmunoResearch, 706-475-148), Alexa Fluor® 594 AffiniPure Donkey Anti-Rabbit 1:500 (Jackson ImmunoResearch, 711-585-152), Alexa Fluor® 647 AffiniPure Donkey Anti-Mouse 1:250 (Jackson ImmunoResearch, 715-605-151). Between each step, the sections were washed with TBS at room temperature. Finally, the sections were mounted with 200 μl Prolong™ diamond antifade mounting solution (Invitrogen, P36961) and covered with CoverWell™ incubation chamber gasket (Life Technologies, C18150).

### Automated serial section imaging

For light microscopy, we used a motorized inverted microscope, Eclipse Ti-E (Nikon, Eclipse Ti-E) with 100×, 1.45 N.A., oil-immersion objective (Nikon, Plan APO γ) (Figure 1B_6_). We collected 2 × 3 image tiles with 15% overlap with four color channels (Figure 1B_7_).

### Image refinement and alignment

All images were histogram equalized to enhance image contrast using Metamorph (Miyamoto et al., 2016) to match them with a selected image with high contrast. The 2 × 3 image tiles were montaged by a custom-built macro in Metamorph. The 297 montaged sections were aligned using TrakEM2 (Cardona et al., 2010). Synaptophysin-1 images were used for alignment with manual correction as required.

### Dendrite tracing and synapse annotation

For tracing dendrites, we alternately used Omni (Kim et al., 2014), pyKnossos (Wanner et al., 2016), and TrakEM2. Custom-built MATLAB codes were used to freely convert the coordinates among the software. A rough volumetric reconstruction of dendritic branches was obtained by thresholding the aligned GFP image stack and was visualized on Omni by 3D mesh rendering. We selected 18 relatively well conserved dendritic branches based on the mesh image. The exact dendritic structures were manually traced using pyKnossos with the help of Omni. pyKnossos provides multiple “viewports,” including the plane that is perpendicular to the process under tracing. The nodes were placed sparsely for straight parts of the branches and more frequently at the locations of sudden changes such as branching points or highly curved parts. The intervals between adjacent nodes vary roughly from 0.2 μm to 1 μm. The multiple viewports and 3D skeleton representation on pyKnossos ensured rapid and accurate tracing. The possibility of reconstruction errors was further investigated by comparison of the skeleton representation on pyKnossos to the volumetric representation of the putative neurons on Omni. By the comparison, it is readily noticed when the tracing trespasses into a different neuron and makes a merger or when the tracing fails to continue in existing branches and makes a split.

Finally, the locations of synapses with the two origins were annotated using TrakEM2, those from one input origin at a time. Three channels of the fluorescence images were displayed for each synapse type. Using the RGB colors, the locations were annotated as synapses where the dendritic spines are contacted by the axonal processes of interest with synaptophysin-1 staining. The use of synaptophysin-1 gradient together with the axon labeling as an indication for the direction of the synapse enhanced the accuracy of synapse detection (Figure 2B and C).

### Computational analysis

The synapse distribution pattern was computationally analyzed from the synapse-tagged skeletons using in-house written MATLAB codes. For the ease of data handling, the skeletons were converted from the node-connected form to a voxel-series form before the analyses as follows. As a preprocess, all the synapse- and non-synapse nodes were labeled with a tag for the branchlet they belong to. This label was used later to assign the synapses to their associated branchlet. Then, the non-synapse skeleton nodes and synapse-tagged ones are processed separately. First, the non-synapse nodes of the skeletons were allocated to corresponding voxels. The voxels are dilated by morphological operation until they form a single connected component. This volumetric branch was thinned again into a voxel-series skeleton by an algorithm that finds the central path of 3D objects (M. Sato. et al., 2000). Finally, the synapse-tagged nodes are projected onto their respective closest skeleton voxel belonging to the same branchlet.

1. **Shuffling of synapses.** In various analyses throughout the manuscript, synapse-shuffled dendritic branches were used as a control group relative to the experimental group of the observed dendritic branches. In the control group, the synapses on dendritic branches exist at random locations while the structures of the branches remain the same as the experimental group. To locate the synapses randomly, synapse shuffling was performed by Monte Carlo simulation that randomly chooses *N_i_* separate synapse locations among *M_i_* skeleton voxels, where *N_i_* is the number of synapses and *M_i_* is the number of voxels of dendritic branch *i*. Different numbers of synapse-shuffled dendritic branches were generated for different analyses.
2. **Dendritic tropism.** The tropism is a tendency that the synapses with the same origin are clustered on a branchlet. The number of synapses and the linear synapse density versus the branchlet length were analyzed. The number of synapses in a branchlet was counted using the branchlet tag of the synapse. The length of a branchlet was calculated by the sum of the path length of the skeleton voxels. To statistically test if there exists a correlation between the linear synapse density and branchlet length, a permutation test based on Pearson’s correlation coefficient was performed. In addition, the preference index was introduced to determine whether there exists a preference in a branchlet depending on the input source of the synapses. The preference index of a branchlet *i*, *Pl_i_*, is defined as follows:

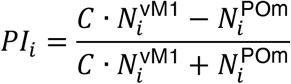

where 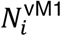 and 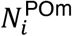 are the number of synapses in the branchlet *i* from vM1 and POm, respectively, and 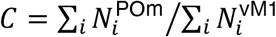 is the normalization factor to compensate for the difference in the total numbers of POm and vM1 synapses in a branch. Similar quantities were measured for terminal branchlets. The terminal branchlets were identified as those containing the end skeleton voxel with no further continuation. The pia distance of synapse was approximated by the distance from the top of the image stack along the z-axis.
3. **Cluster analysis.** We used a single-linkage, unsupervised,hierarchical clustering method using the mean nearest neighbor distance as a dissimilarity criterion. First, the distance between pairwise synapses was calculated by 3D path length along the dendrite. Then, the nearest synapses were grouped together when the distance between adjacent synapses was shorter than the mean value of the nearest neighbor distance. The grouping continued sequentially to a higher level. Intuitively, synaptic clusters should have three or more synapses, with the density higher than the mean synaptic density of the branch. Therefore, we iteratively pruned the outermost synapses if the group of synapses has a density lower than the mean synapse density. If the remaining group has more than three synapses, we finally determined the group as a synaptic cluster (Yadav et al., 2012). We followed the C-score concept based on Monte-Carlo simulation to distinguish highly clustered branches and sparse clustered branches (Yadav et al., 2012). This concept referred to the ratio of the simulated branches with less synaptic clusters than the branch of interest to all the simulated dendritic branches. To generate the simulated branch in this study, the positions of synapses were randomly changed 10,000 times while maintaining the total number of synapses. Consequently, the higher C-score a branch has, the greater fraction of the random branches has a smaller number of synaptic clusters than that of the observed branch. Based on this value, we confirmed the existence of highly clustered branches among the branches we reconstructed. This was expressed in the formula as follows:

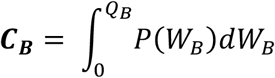 In this, W_B_ is the number of clustered spines in the simulated branches, *P*(*W_B_*) is the probability distribution, and *Q_B_* is the number of clustered synapses in the measured branch for which we want to calculate a C-score. Also, we performed a binomial test to examine the hypothesis that the number of highly clustered branches would exist more than randomly expected (Figure 5E-F). The *P* value derived from the binomial test indicates the probability that more highly clustered branches than observed (m-1) can be found by chance. When the total number of the branch is [n] and the observed number of highly clustered branches [*m* – 1], the *P* value can be calculated from the following formula.

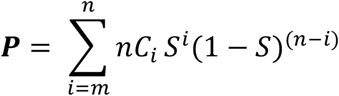

*nC_i_* is the number of combinations when the total number of branches is [*n*] and the number of highly clustered branches is [*i*]. *S* is the probability that the branch is highly clustered above a given threshold, which equals (1 – C-score). To analyze the distance between clusters of different origins, we measured the centroid distance between each POm cluster and the nearest vM1 cluster. For comparison with a randomized control, we created 100 simulated branches by randomly changing the locations of the clusters, while maintaining the total number and characteristics of clusters of vM1 and POm.
4. **Synaptic cloud density analysis.** To quantify the proximity of the clusters by two synapse types, we introduced the synaptic density cloud model. In this model, a synapse is regarded to exist probabilistically. The spatial probability density function for the location of a synapse *i*, that is the probability to find the synapse *i* at the position *x* on a single path of a dendrite, follows the Gaussian distribution as below:

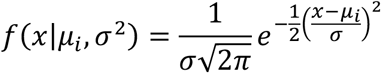

where mean *μ_i_* is the actual location of the synapse and standard deviation *σ* is a given value. Mathematically, when the standard deviation goes to zero, the Gaussian distribution function becomes Dirac’s delta function. This means that the synapse is located at its actual location with probability 1 and the deterministic synapse location is restored. Here, we heuristically chose the value *σ* = 500 nm, which makes the Gaussian curve decay reasonably in the scale of the dendritic branches. On each bifurcation of dendrites, the spatial probability density was divided into halves. Therefore, the probability to find the synapse *i* at the position *x* on the dendritic branch becomes as follows:

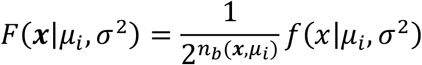 Where *n_b_*(*x,μ_i_*) is the number of bifurcations on the path from *μ_i_* to *x*. Note that now *x* represents an arbitrary location on the arborized dendritic branches instead of on a single path. The synaptic cloud density for an input type is defined as the sum of all the synapses of the type:

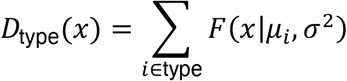 Theoretically, this is a continuous function of *x*, but we discretized this quantity by taking the bin average over every 100 nm intervals in this study.

### Statistical Analysis

The Kolmogorov-Smirnov test was used to determine nearest neighbor distances between the synaptic inputs from vM1 or POm in comparison to the random distribution. It was also used to compare distances to centroids of the nearest vM1 synaptic cluster from the centroid of the POm synaptic cluster and the distances between randomized synaptic clusters. One-way ANOVA was used to compare C-score distribution from branches with measured synapses and separately simulated 540 random synaptic locations.

The one-sample t-test was used to make a comparison between the number of branches with highly clustered synapses from vM1 and POm and the same set of branches with randomized synaptic locations. Wilcoxon matched-pairs signed-rank test was used to determine the fraction of synapses participating in synaptic clusters of vM1 and POm and the cluster occupancy. The Mann Whitney test was used to validate the number of synapses in a vM1 and a POm cluster and the integrated cluster length for vM1 and POm synapses. The Wilcoxon rank sum statistic was applied for testing the distribution of SCD_vM1_ on the dendritic fraction with high SCD_POm_ compared with low SCD_POm_.

## ACKNOWLEDGMENTS

Supported by grants from the KBRI Research Program (20-BR-01-01 and 20-BR-04-01) and Brain Research Program through the National Research Foundation of Korea (NRF) of the Ministry of Science and ICT (NRF-2017M3C7A1048086 and No.2017M3A9G8084463). All the fluorescent imaging data were acquired at the Advanced Neural Imaging Center of KBRI.

## AUTHOR CONTRIBUTIONS

J-CR designed the experiment with input from JHC and NK as well as others in the Rah laboratory. NK collected the data and NK, SB and JHC analyzed the data with the supervision of JSK and J-CR. JSK and J-CR interpreted the data and wrote the manuscript with input from the other authors.

## DATA AND CODE AVAILABILITY

All data supporting the findings of this study are provided in the main text and SI Appendix. All the codes for the data analyses are available at https://github.com/cns-kim-lab/nkim_et_al_2020. The annotation data containing skeleton reconstruction with synapse tags can be found with the codes. The aligned 3D images are available upon request.

## SUPPLEMENTAL INFORMATION

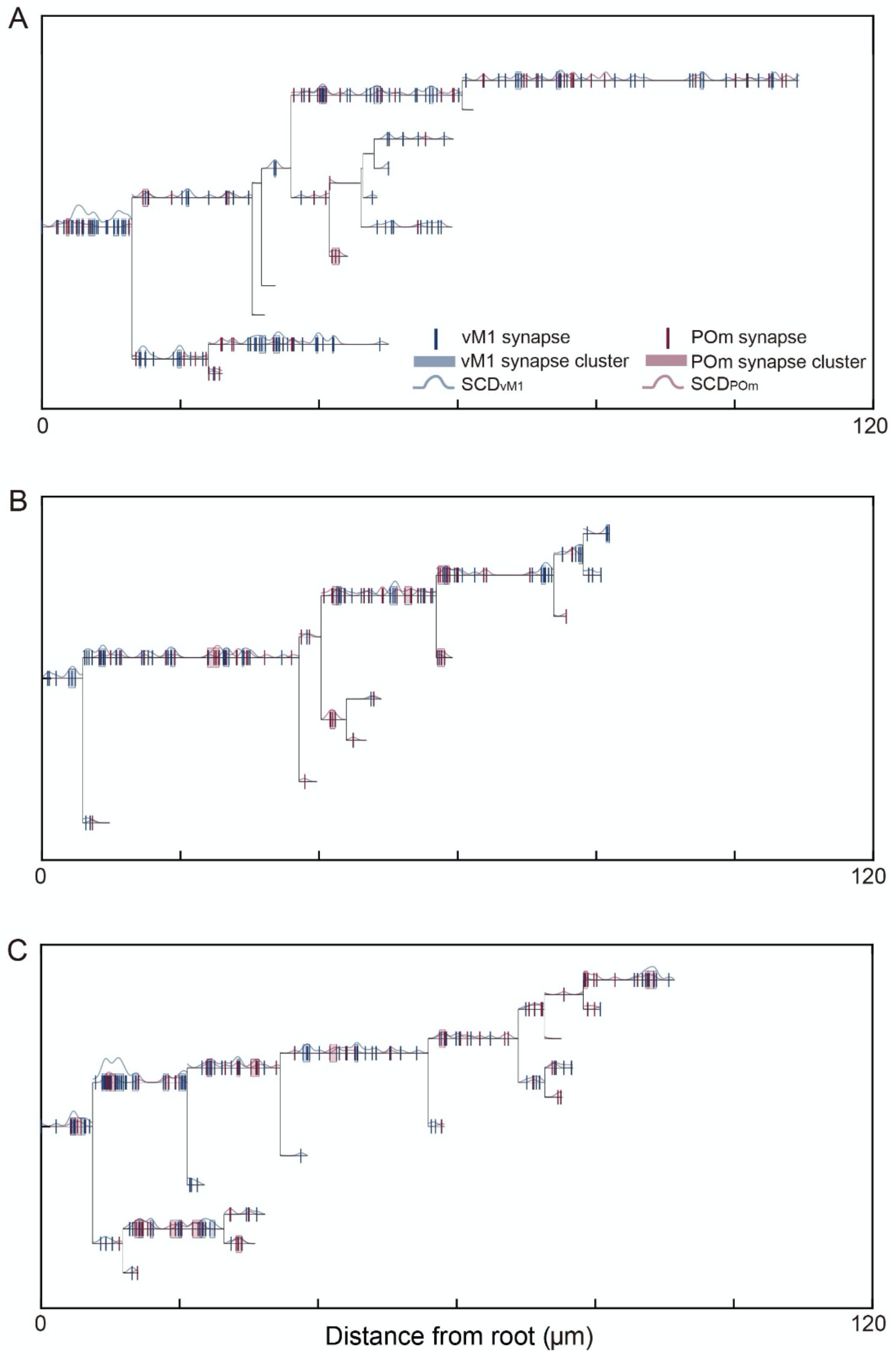
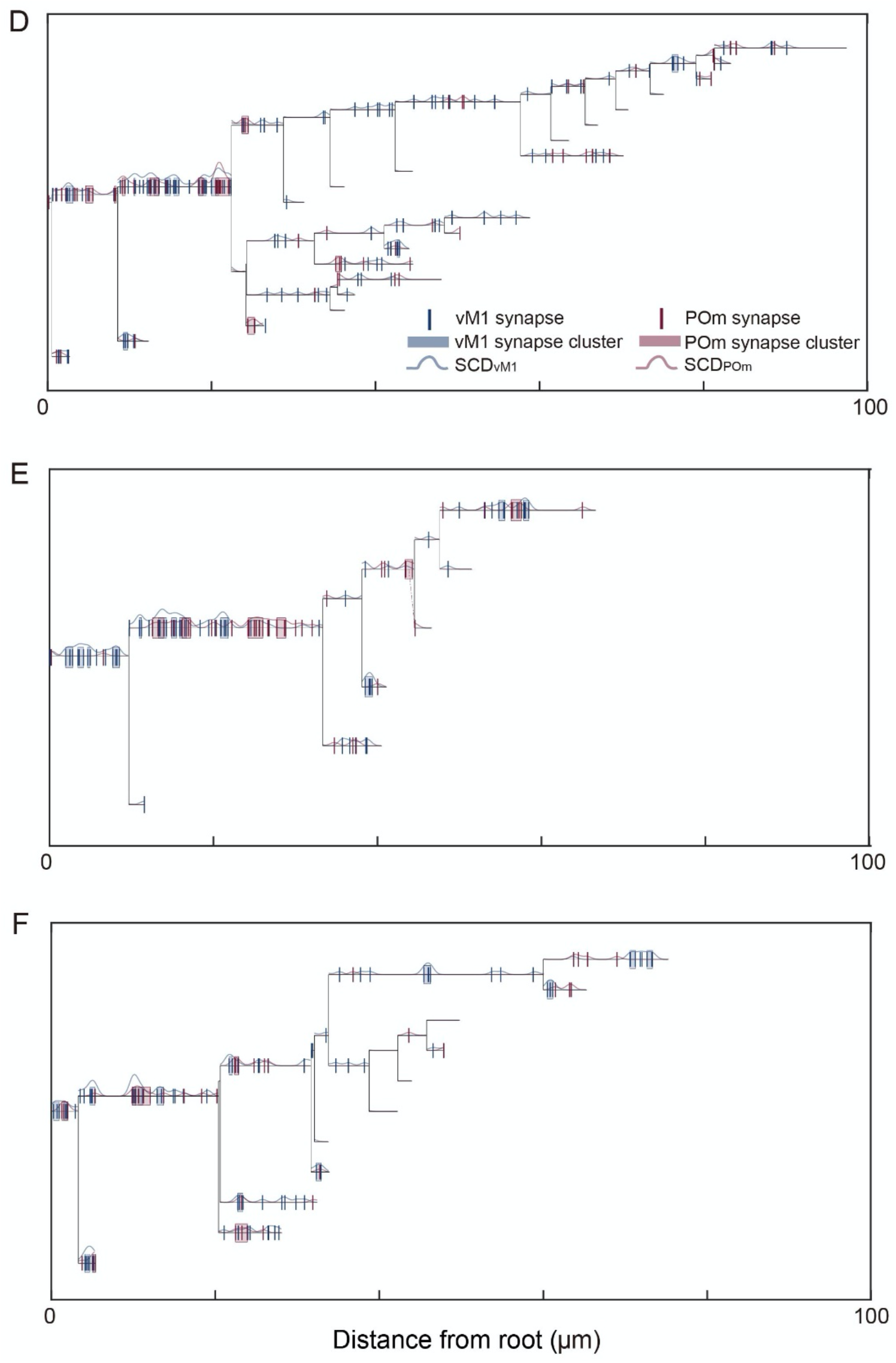
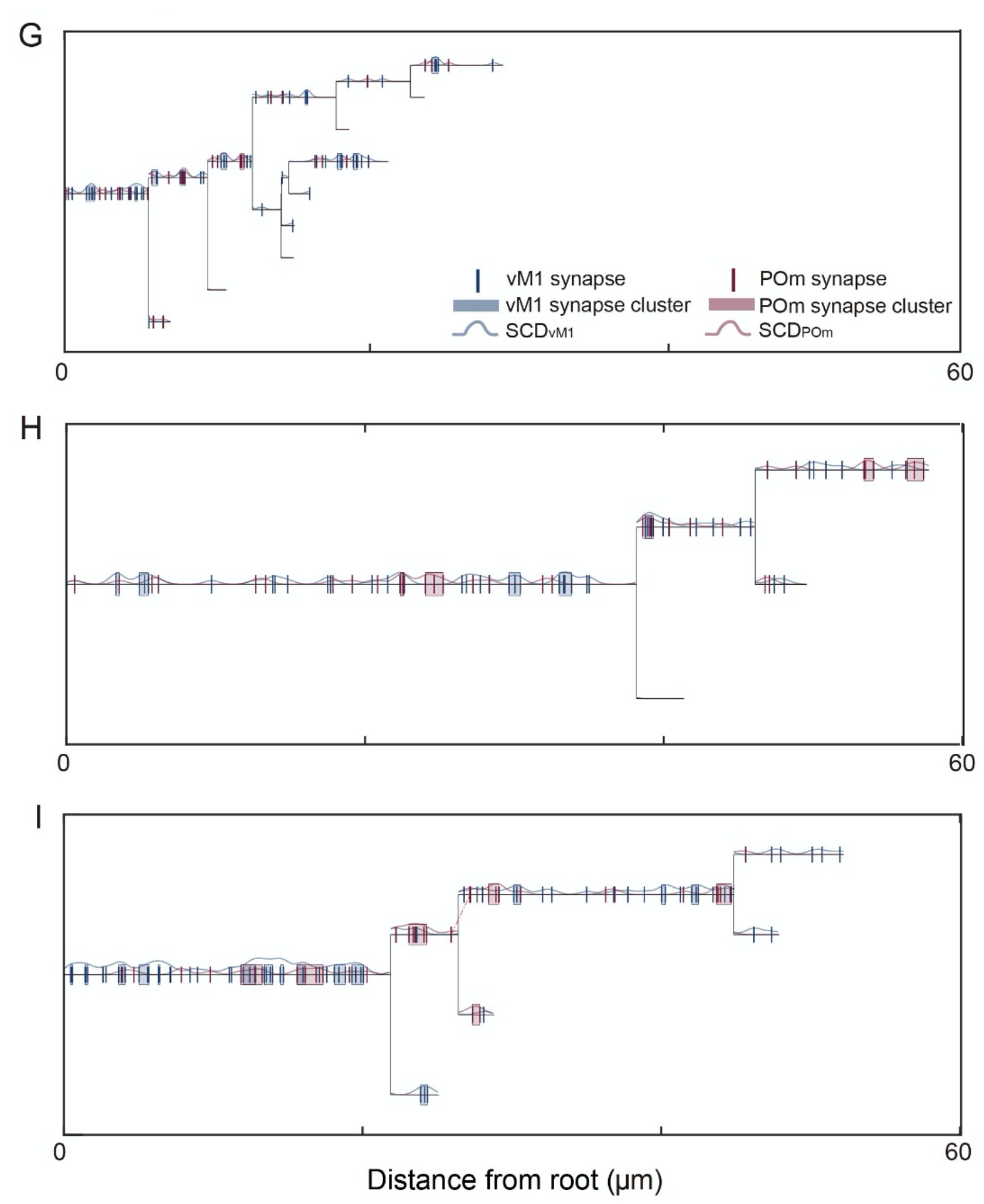
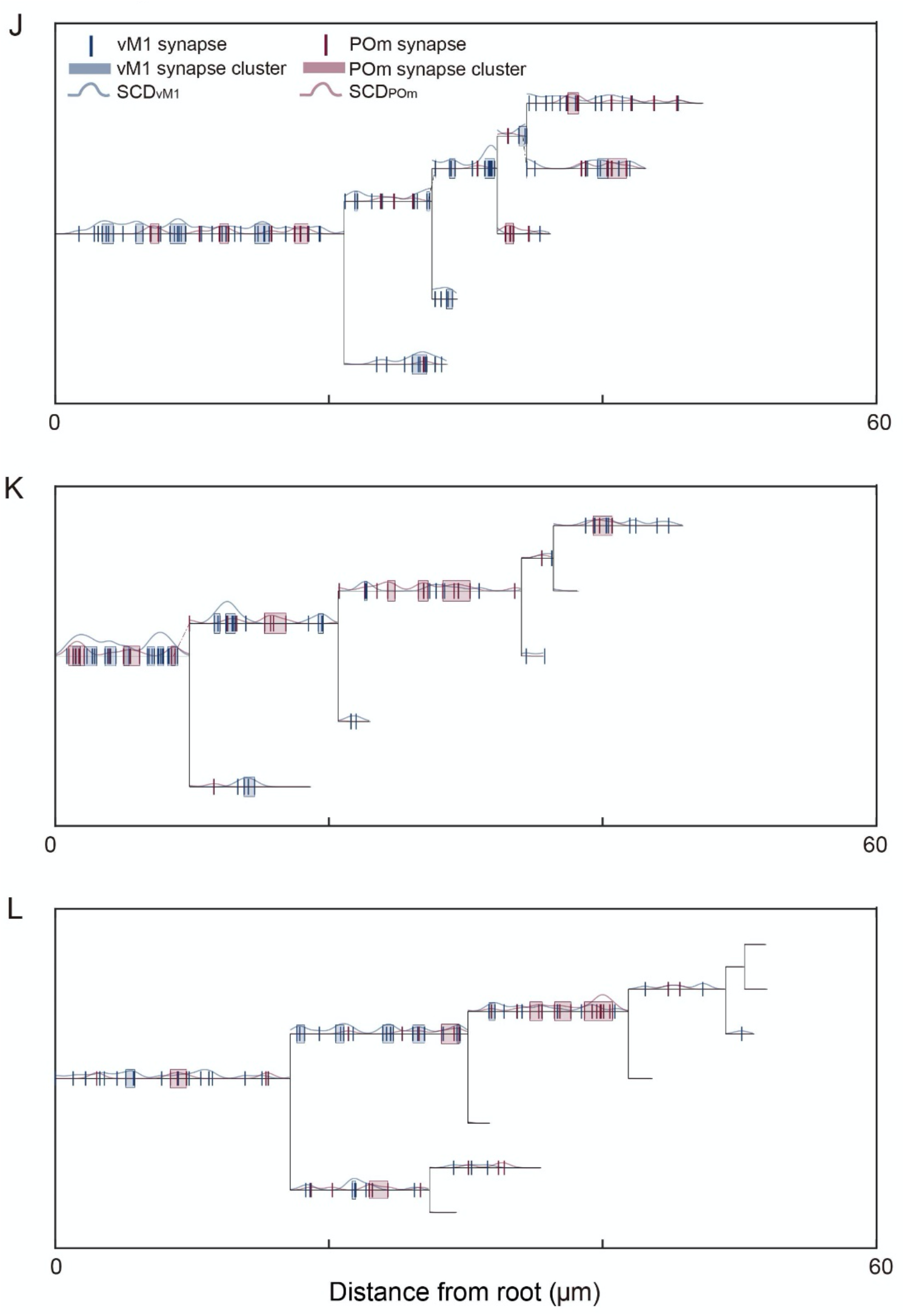
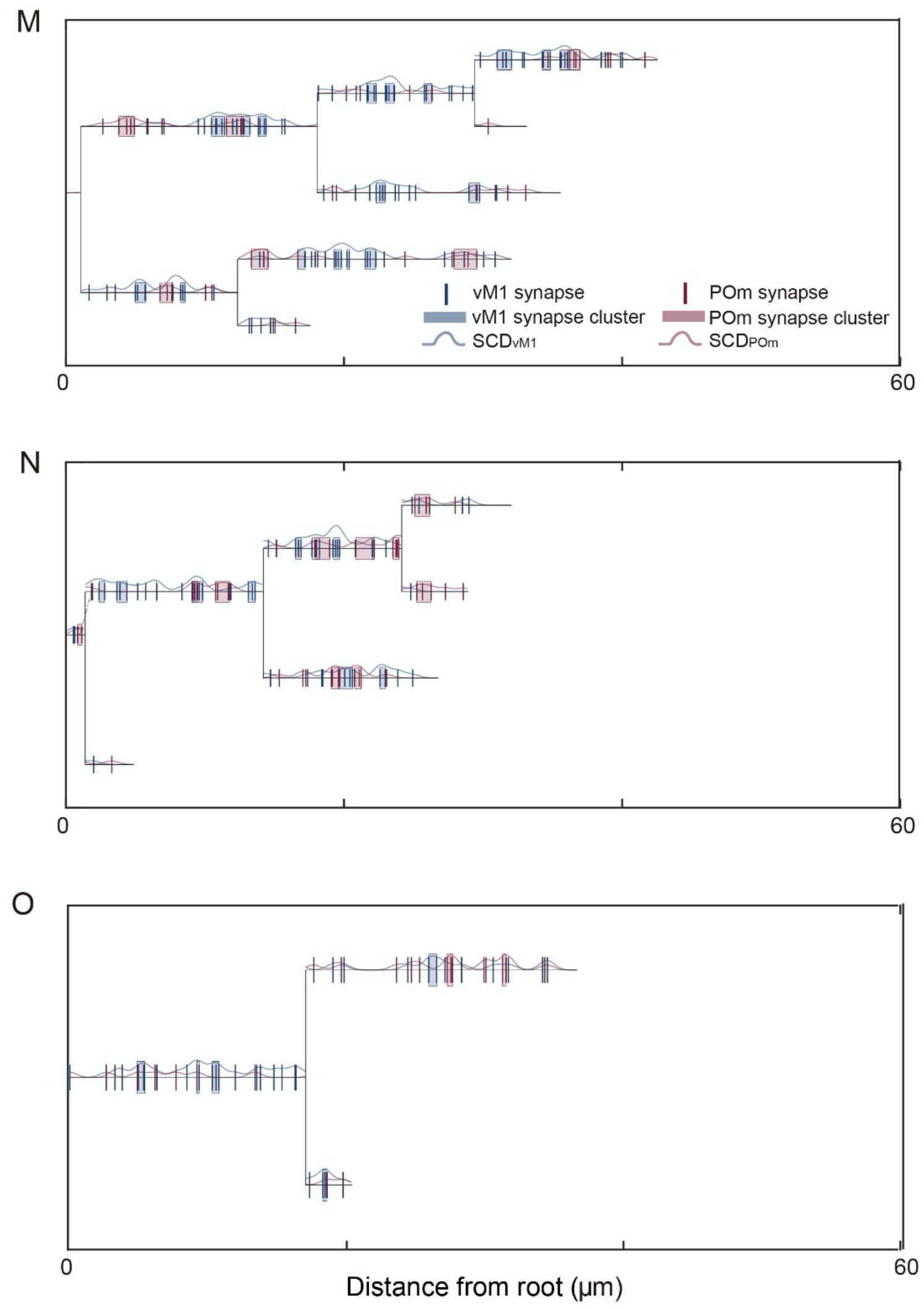
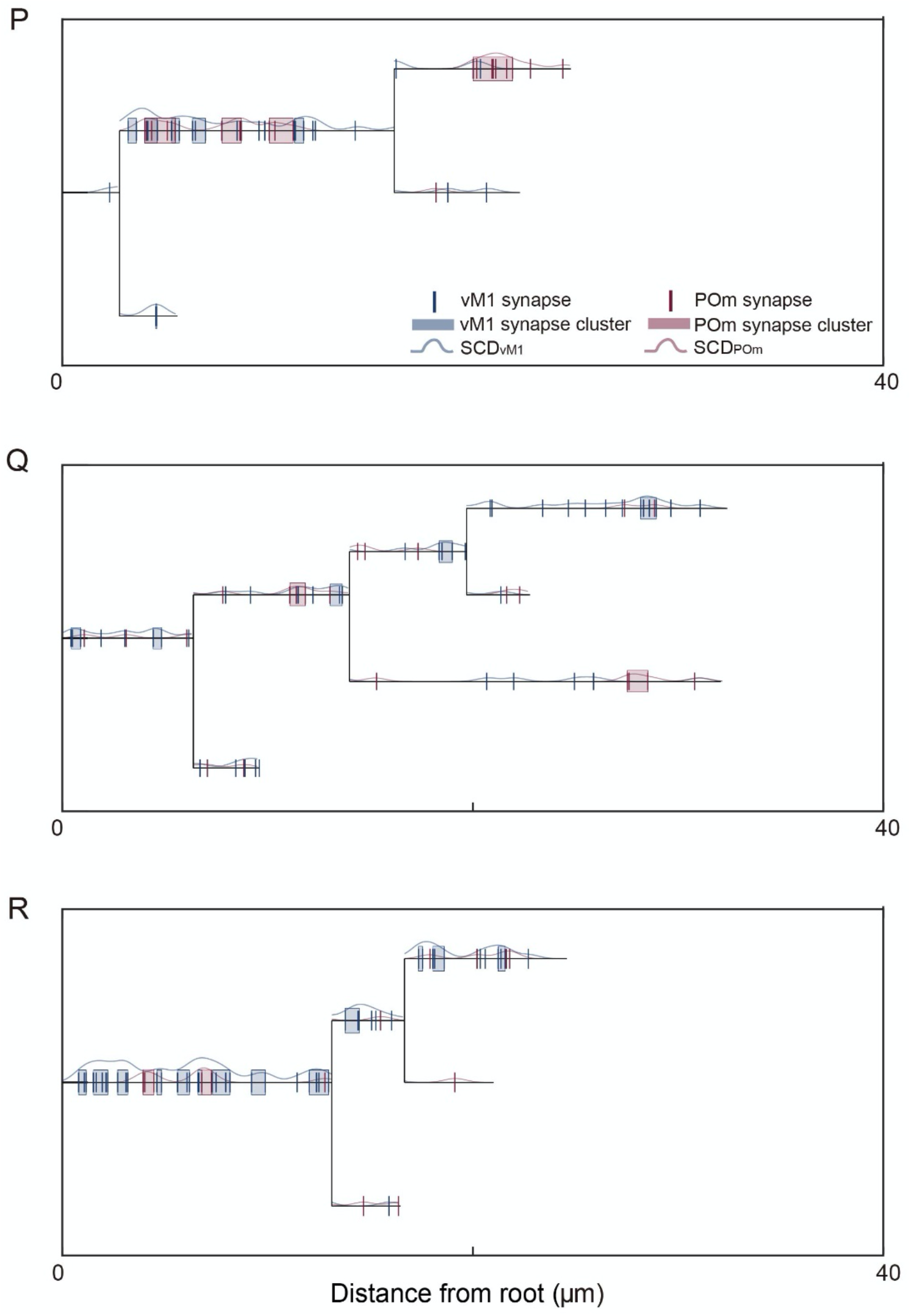

**Related to figure 5 and 8.** Dendrograms from all the reconstructed dendrites with the synaptic cloud density and the location of synapses from vM1 (blue) and POm (red). Groups of synapses determined as synaptic clusters were boxed in blue (for vM1 synapses) or in red (for POm synapses).

